# Hypoxia induces early neurogenesis in human fetal neural stem cells by activating the WNT pathway

**DOI:** 10.1101/2021.07.14.452315

**Authors:** D Dey, D Joshi, V Shrivastava, CMS Singal, S Tyagi, MA Bhat, P Jaiswal, JB Sharma, JK Palanichamy, S Sinha, P Seth, S Sen

## Abstract

Fetal neural stem cells (FNSCs) are present in the brain of human fetuses that differentiate into cells of neuronal and glial lineages. Difference in oxygen concentration between maternal and fetal circulation indicates that the developing fetus may be exposed to lower oxygen concentrations compared to adults. This physiological hypoxia may influence the growth and differentiation of the FNSCs. This study aimed to evaluate the effect of hypoxia on the differentiation potential of human FNSCs isolated from the sub-ventricular zone of aborted fetal brains (n=5). FNSCs were isolated, expanded, and characterized by Nestin and Sox2 expression, using immunocytochemistry and flowcytometry respectively. These FNSCs were exposed to 20% oxygen (normoxia) and 0.2% oxygen (hypoxia) concentrations for 48 hours, and hypoxia exposure (n=5) was evaluated by a panel of markers (CA9, PGK1 and VEGF). Whole human genome transcriptomic analyses (Genespring GX13) of FNSCs exposed to hypoxia (Agilent 4×44K human array slides), highlighted that genes associated with neurogenesis were getting enriched. The pathway analysis of these enriched genes (using Metacore) showed that WNT signaling played a role in determining the cell fate of FNSCs exposed to hypoxic environment. Microarray analyses was validated using neuronal and glial lineage commitment markers such as Ngn1, Ngn2, ASCL1, DCX, GFAP, Olig2 and Nkx2.2 using qPCR (n=9). This demonstrated upregulation of the neuronal commitment markers on hypoxia exposure, while no change was observed in astrocytic and oligodendrocyte lineage commitment markers. Increased expression of downstream targets of the WNT signaling pathway, TCF4 and ID2, by qPCR (n=9), indicated its involvement in mediating neuronal differentiation on exposure to hypoxia.

## Introduction

Neural stem cells (NSC) are multipotent cells that can differentiate into neurons, astrocytes and oligodendrocytes. Fetal neural stem cells (FNSCs) are located in the subventricular zone and dentate gyrus of the fetal brain. NSC are also found in the cortex, striatum and subependymal zone of adult brain (Kjell et al., 2020; Bond et al., 2015; Yin et al., 2013). In addition to cues provided by the NSC niche, oxygen concentration can also influence the growth and differentiation potential of NSCs, and plays a critical role during embryonic development (Vecera et al, 2020; Qi et al, 2017). As the fetus develops inside the uterus, the difference in oxygen concentration between maternal and fetal circulation shows that the developing fetus is normally exposed to lower oxygen concentrations, and thus, despite fetal hemoglobin having a greater affinity for oxygen, there is a possibility that the fetus is exposed to hypoxia *in utero* (Cummings et al, 2017; Martin et al., 2010). Studies have shown that hypoxia may influence NSC development and plasticity (Qi et al, 2017; De Filippis and Delia, 2011). It has been reported that mild hypoxia (5% O_2_) activates molecular pathways like Wnt/beta-catenin and Notch, which regulate self-renewal and proliferation of stem cells, including NSCs (Sun et al., 2021; Qi et al, 2017; Felfly et al, 2011).

This study aimed to understand the role of hypoxia on differentiation potential of human FNSCs. It also elucidates the possible mechanism by which hypoxia may influence lineage commitment in human FNSCs.

## Methodology

### Sample collection

Aborted fetal samples were collected from the Department of Obstetrics and Gynecology, AIIMS, New Delhi, India. Informed consent was obtained from mothers undergoing Medical Termination of Pregnancy (MTP) in their second trimester of pregnancy (12-20 weeks) for maternal indications. Mothers undergoing MTP for fetal indications (such as chromosomal anomalies) were excluded from the study. Approvals were taken from Institutional Ethics Committee and Institutional Committee for Stem Cell Research, before starting the study. The study was carried out in conformation with the Helsinki Declaration.

### Isolation of human fetal neural stem cells (FNSCs)

Isolation of human FNSCs from the brain of aborted fetuses was done as per published protocol (Bhagat *et al.*, 2018). Briefly, tissue from subventricular zone of brain was isolated and plated onto poly-D-lysine coated culture flasks in neural stem cell media containing neurobasal media (GIBCO, NY, USA) with 1% N2 supplement (GIBCO, NY, USA), 2% Neural survival factor-1 (Lonza, IA, USA), 1% Glutamax (GIBCO, NY, USA), 5mg/mL of bovine serum albumin (Sigma, MO, USA), penicillin (50 IU/ml), streptomycin (50 μg/ml) and gentamicin (2 μg/ml). Tissue demonstrating cells radiating from the core, were gently dissociated and subcultured onto poly-D-lysine coated flasks, to generate monolayers of FNSCs.

### Flow cytometry

Human FNSCs were fixed with 2% paraformaldehyde, permeabilized with 1% BSA containing 0.1% Triton X-100. Cells were blocked with 2% BSA for half an hour and subsequently stained with (intracytoplasmic) mouse anti-human SOX2 antibody conjugated with V450 (BD Biosciences, cat. no. 561610) using appropriate controls. Cells were washed, resuspended in 2% paraformaldehyde, and data was acquired using BD LSR Fortessa (BD Biosciences, San Jose, CA, USA) and analyzed using FlowJo v10 software.

### Immunocytochemistry

Human FNSCs were plated onto coverslips coated with poly-D-Lysine. They were washed with PBS and fixed with 2% PFA. Cells were incubated for 1 hour in blocking solution (1% BSA with 0.1% Triton X-100) and then washed with PBS. The cells were incubated overnight at 4°C with primary antibody (Mouse anti-Nestin 1:1000, Cat no. 33475, CST, MA, USA; Rabbit anti-SOX2 1:1000, Cat no. 23064, CST, MA, USA). The cells were washed thrice with PBS and then incubated with secondary antibody (Mouse anti-Rabbit FITC, 1:1000: Cat no. A11008, Invitrogen; Goat anti-Mouse Alexa Fluor 594, Cat no. A-11005, Invitrogen) for 1 hour at room temperature. Cells were then washed thrice with PBS, and mounted onto glass slides using Vectashield mountant containing DAPI. The slide was allowed to dry overnight. Images were taken on Nikon Eclipse Ti-S fluorescent microscope (Tokyo, Japan) and analyzed with NIS-Elements BR software.

### Exposure of human FNSCs to different oxygen concentrations

Human FNSCs were exposed to oxygen concentrations mimicking normoxia (20% oxygen) and hypoxia (0.2% oxygen) for 48 hours, at 37°C and 5% CO_2_, that was created using an Anoxomat hypoxia induction system (Advanced Instruments, Norwood, MA, USA). Hypoxia exposure was validated by evaluating the expression of hypoxia-responsive genes CA9, VEGF and PGK-1.

### RNA isolation, cDNA synthesis and qPCR

Total RNA was extracted from the cells after exposure to different oxygen concentrations, using Tri-Reagent (Sigma, MO, USA) and quantified by Nano-Drop ND-1000 spectrophotometer (Thermo-Fisher Scientific, MA, USA). cDNA was synthesized with 1μg total RNA using M-MuLV-RT (Thermo-Fisher Scientific, MA, USA) and random hexamer primers (IDT, IL, USA). The expression of various genes was evaluated in the cells (in triplicates) using gene-specific primers (IDT, IL, USA) (Table 1) and DyNAmo Flash SYBR Green qPCR kit (Thermo-Fisher Scientific, MA, USA) using CFX96 Touch™ Real-Time PCR Detection System (BioRad, CA, USA). 18S rRNA was used as an internal reference gene for normalization. Relative fold change in gene expression was calculated using 2^−ΔΔCT^ method. Human FNSCs exposed to normoxia (20% oxygen) were used as controls for hypoxia experiments.

**Table 1:**
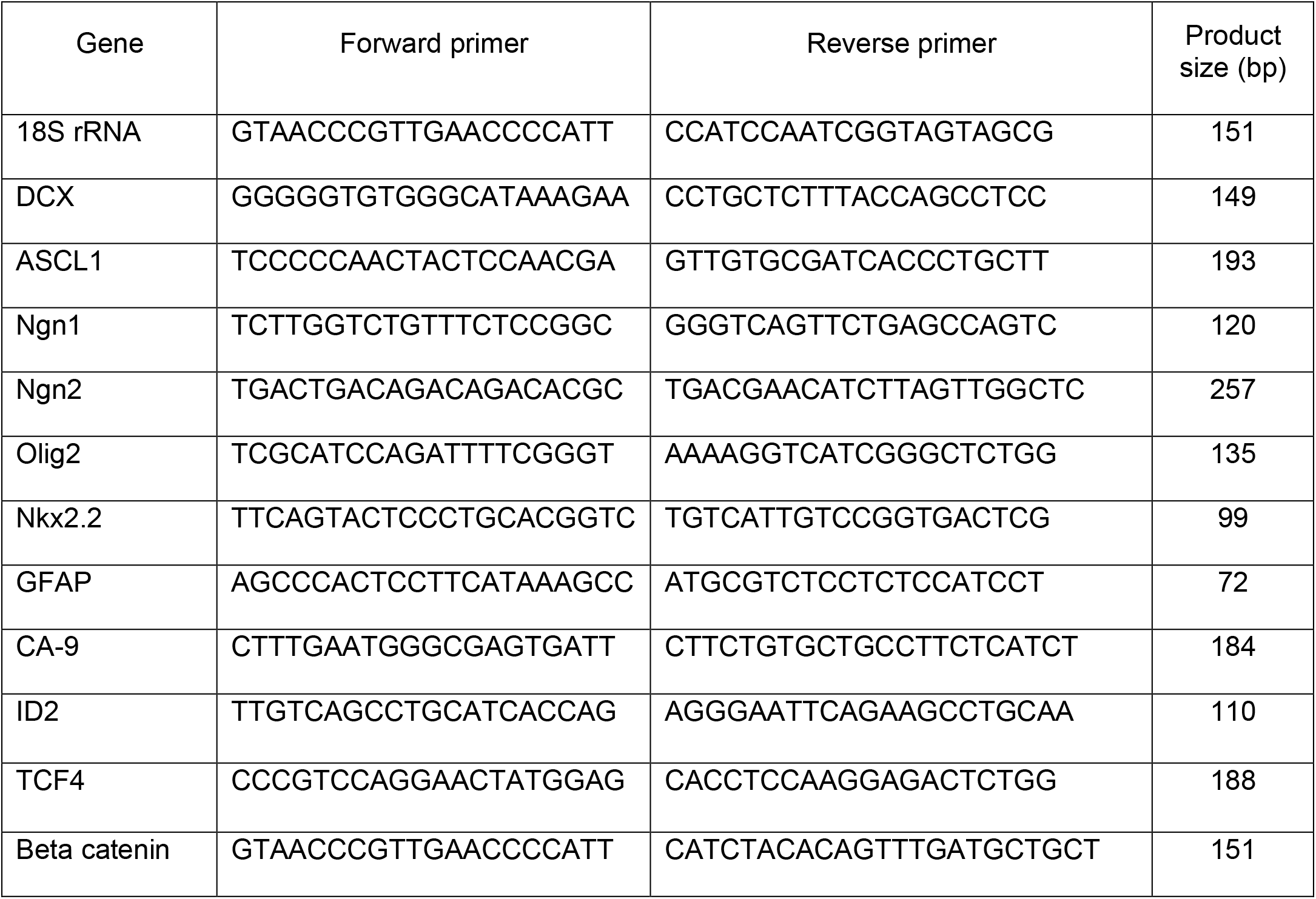
Sequence of primers used for gene expression analyses

### Gene expression microarrays

RNA concentration and integrity of the cells were checked using Nanodrop (Thermo-Fisher Scientific, MA, USA) and Bioanalyzer (Agilent, Santa Clara, CA, USA) respectively. Expression microarray was done on two biological replicates each, of human FNSCs exposed to normoxia and hypoxia, on 4 × 44K human expression array slides (G2519F) (Agilent, Santa Clara, CA, USA), following manufacturer’s protocol. Briefly, a total of 200 ng total RNA per sample (n=2) were subjected to cDNA conversion and linear amplification with fluorescent labeling for preparing Cy3 labeled cRNA. Complementary hybridization (17 hours at 65 °C) was followed by washing. Slides were scanned at 3 μm resolution, followed by feature extraction using feature extraction software version 10.7.1.1 (Agilent technologies, Santa Clara, USA). Analysis was done using Genespring Software v14.9.1 (Agilent technologies, Santa Clara, USA). Principal component analysis and Hierarchical clustering analysis were done to identify gene target distribution, that showed a consistent difference in expression between normoxia and hypoxia exposed cells. This was followed by differential gene expression analysis, which was done using unpaired t-test using Genespring software (GX v14.9.1). The list of differentially expressed genes were imported into gene onotolgy consortium and Metacore for gene ontology and pathway analysis.

### Statistical analysis

Statistical analysis was done using Graph Pad Prism v6. Statistical differences between normoxia and hypoxia exposed groups was estimated using Mann Whitney test. p-value < 0.05 was considered statistically significant.

## Results

### Isolation and characterization of human fetal neural stem cells

Human fetal neural stem cells (FNSCs) were observed to be radiating out from the core of tissue isolated from the subventricular zone of the brain (Fig1a). On dissociating and subculturing these, small, unipolar monolayer of human FNSCs were obtained (Fig1b). Neurosphere assay displayed their ability to form neurospheres (Fig1c). Immunocytochemical staining helped characterize the human FNSCs, and demonstrated expression of Nestin and Sox2 (Fig 1d). Flow cytometry indicated that more than 90% of FNSCs expressed Sox2 (Fig1e and f).

**Figure 1.**
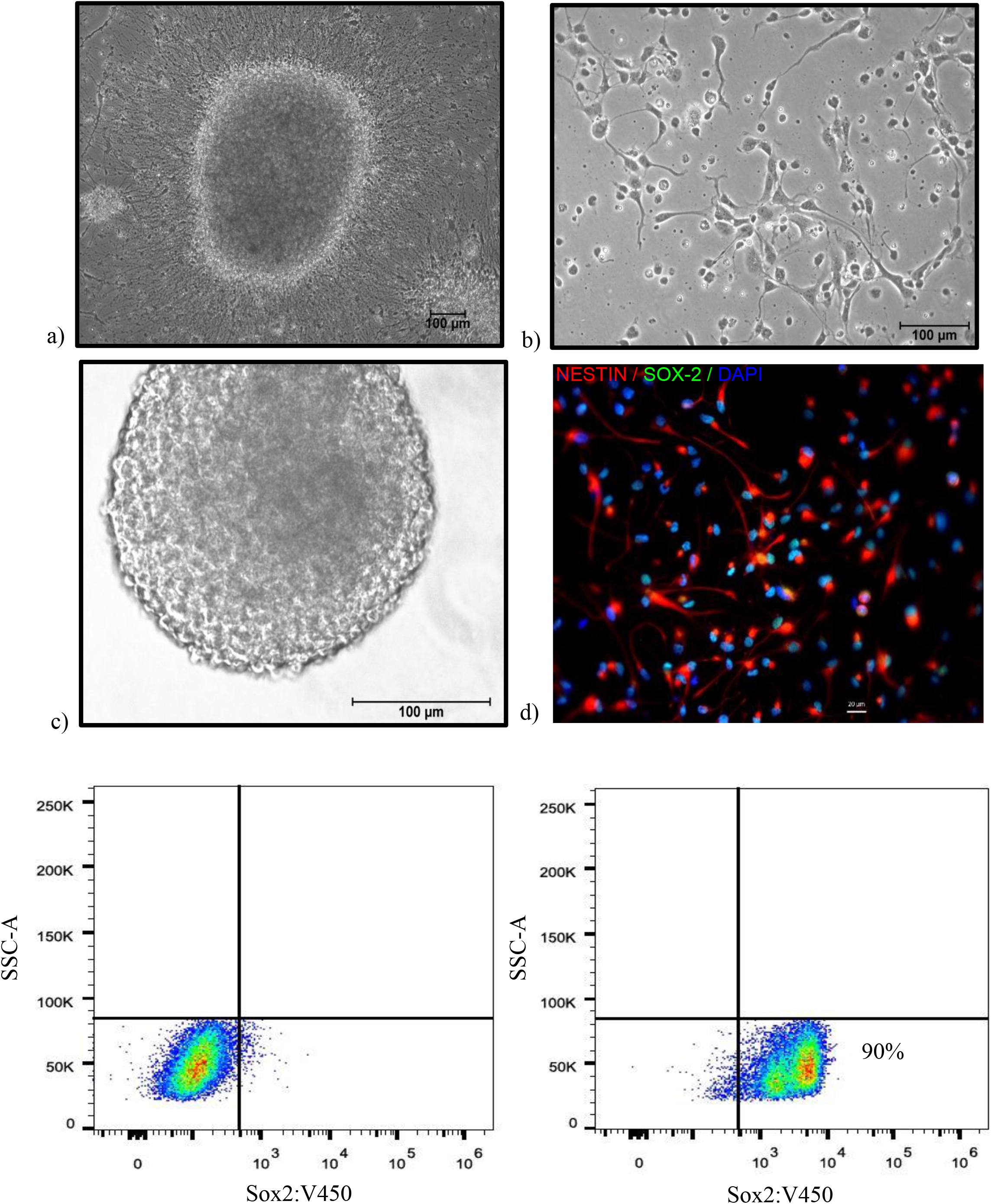
Isolation and characterization of Human FNSCs. a) Human FNSCs radiating out of the core of the tissue isolated from subventricular zone of aborted fetal brain. b) FNSCs in monolayer displaying unipolar morphology. c) Human FNSCs forming neurospheres. d) Immunofluorescence image of FNSCs stained for neural stem cell markers Nestin (red) and Sox2 (green). Nuclei are stained with DAPI (blue). e) Dot plot of unstained human FNSCs in flow cytometry. f) Human FNSCs stained for Sox2 (antibody conjugated with V450).

### Exposure of FNSCs to different oxygen concentrations for 48 hours

Human FNSCs were exposed to different oxygen concentrations mimicking normoxia (20%), and hypoxia (0.2%) for 48 hours. Hypoxia exposure was validated by evaluating CA9, PGK1 and VEGF expression by qPCR. The mean fold change ± SD in CA-9, PGK1 and VEGF expression in the FNSCs exposed to hypoxia were 345.91±38.29 (p<0.0001), 27.61±11.48 (p<0.001) and 6.45±2.56 (p<0.01) respectively (Fig. 2a,b,c).

**Figure 2.**
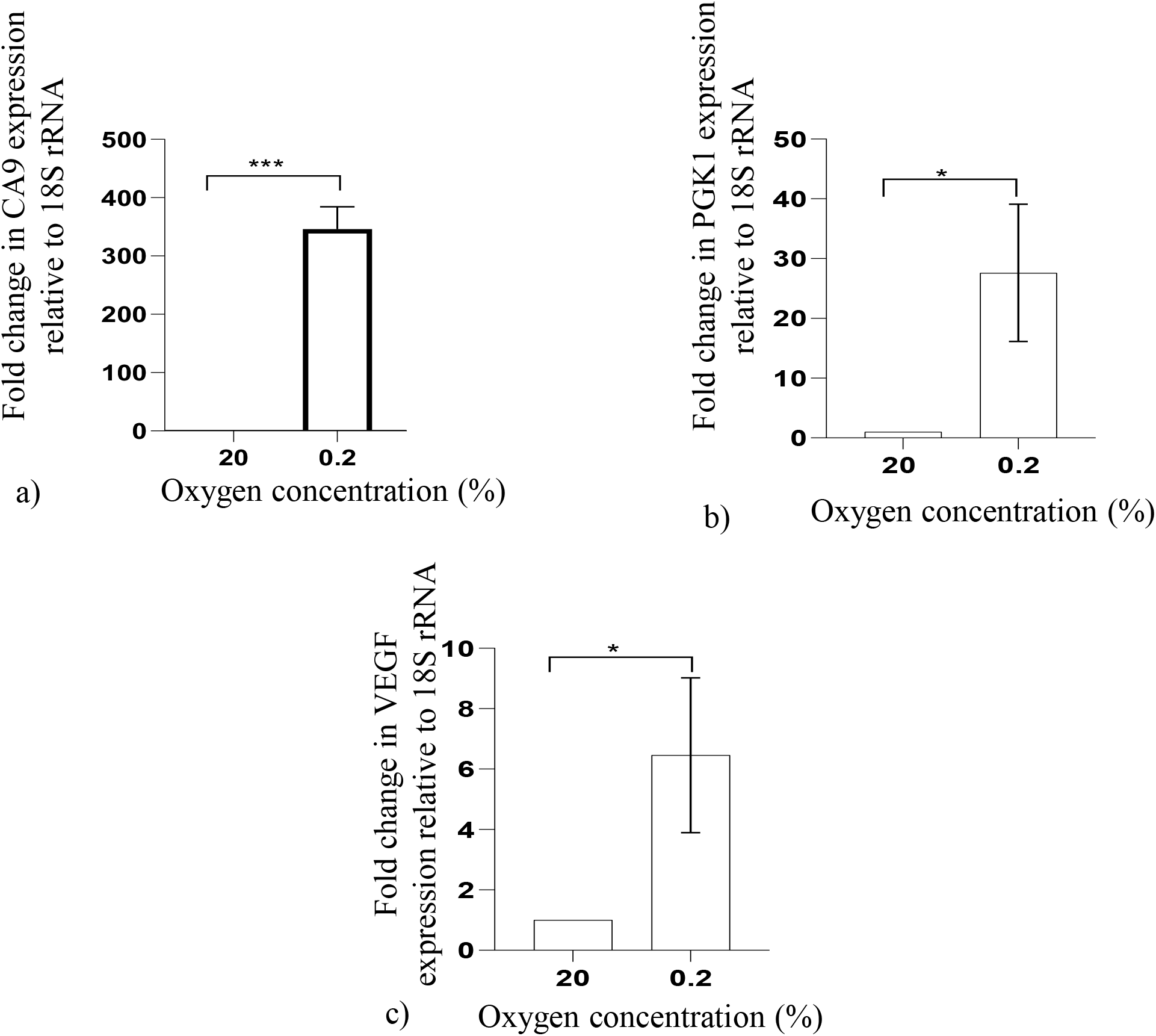
Expression of hypoxia responsive genes, as analysed by qPCR (n=5), when human FNSCs are exposed to hypoxia a) CA9 b) PGK1 and c) VEGF gene expression. Data represented as Mean±SD. * p-value <0.05, ***p-value <0.001.

### Gene ontology and pathway analysis

The differentially expressed genes in FNSCs exposed to hypoxia, when analyzed for gene ontology (GO), showed that cell development and cell differentiation were getting enriched. The genes involved in these GO terms were further analyzed for biological processes, and showed that regulation of neuron projection development, positive regulation of neurogenesis, neuron projection guidance, cell morphogenesis involved in neuron differentiation, regulation of neuron differentiation, neuron projection morphogenesis, neuron projection development, neuron development, and generation of neurons were getting enriched and had role in cell differentiation of human FNSCs exposed to hypoxia (Fig 3a).

**Figure 3.**
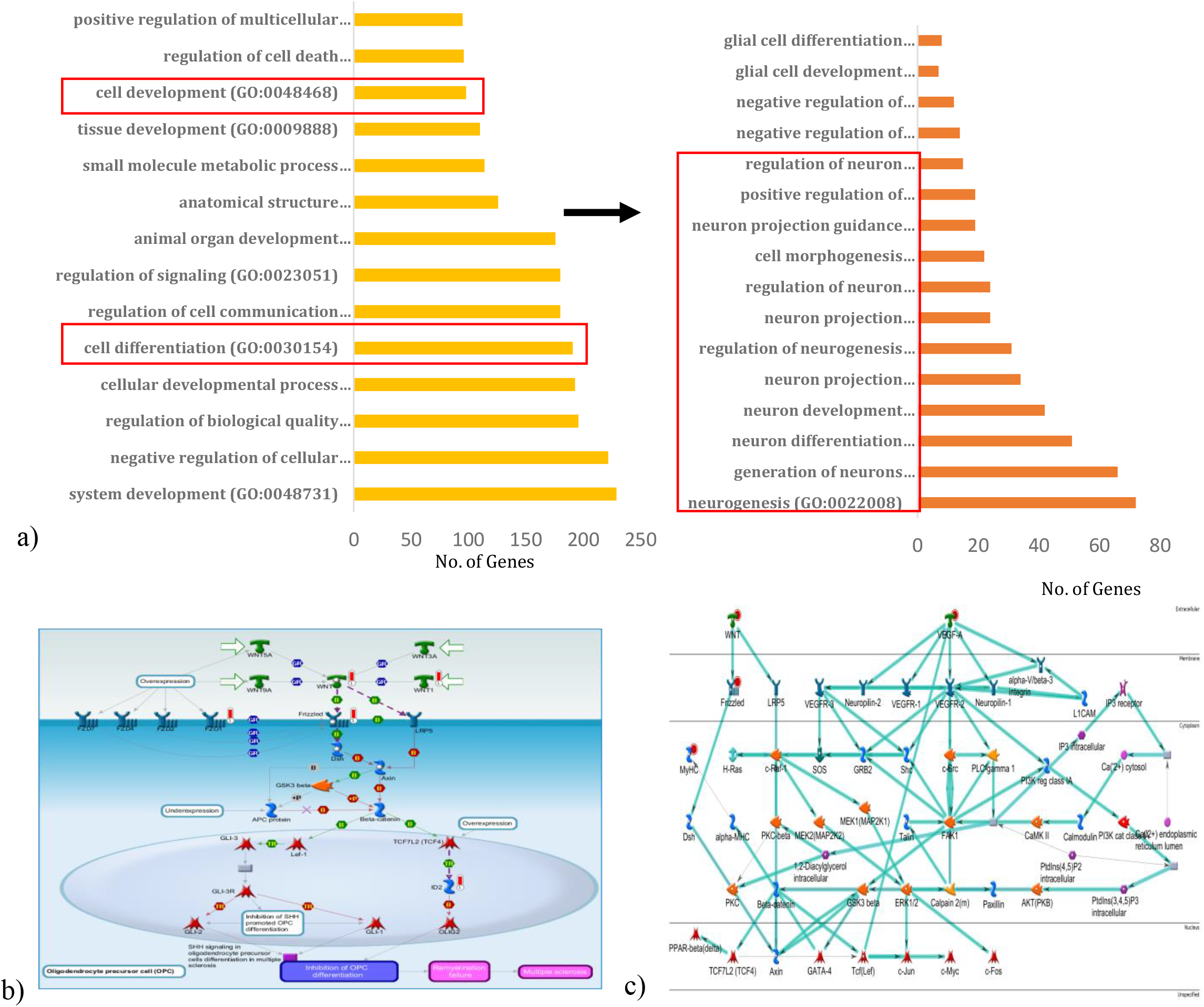
Gene ontology and pathway analysis of human FNSCs exposed to hypoxia (0.2% oxygen). a) Gene Ontology analysis of human FNSCs exposed to hypoxia analyzed in gene ontology consortium. b) Pathway map of genes enriched in neurogenesis after exposure to hypoxia. c) Network map of genes enriched in neurogenesis.

The genes involved in the neuron development, neuron differentiation and generation of neurons were then evaluated for enriched pathways and networks using Metacore software. This analysis showed that Wnt-beta catenin canonical network was involved, with candidate genes such as Wnt and Frizzled being up-regulated (Fig 3b). It was also seen that a sub-node of a VEGF pathway was linked with the Wnt-beta catenin canonical network (Fig 3c).

### Expression of lineage commitment markers in human FNSCs exposed to hypoxia

ASCL1, DCX, Ngn1 and Ngn2 are markers for early neurogenesis, and their expression was checked by qPCR, after exposing human FNSCs to hypoxia for 48 hours. The mean fold change ± SEM for Ngn1, Ngn2, ASCL1, and DCX expression in the FNSCs exposed to hypoxia were 36.08±21.88 (p=0.0035); 0.65± 0.13 (p=0.0035); 3.23± 1.45 (p=0.4042); and 3.08±1.34 (p=0.7090) respectively (Fig 4 a-d). Astrocytic lineage marker, GFAP, and oligodendrocyte lineage markers, Olig2 and Nkx2.2, did not show any significant change when evaluated by qPCR. The mean fold change ± SEM for GFAP expression in the FNSCs exposed to hypoxia, was 1.02±0.20 (p=0.6818) (Fig. 5), while for Olig2 and Nkx2.2, it was 0.98± 0.38 (p=0.2235) and 1.23 ± 0.30 (p=0.7059) respectively (Fig. 6 a,b).

**Figure 4.**
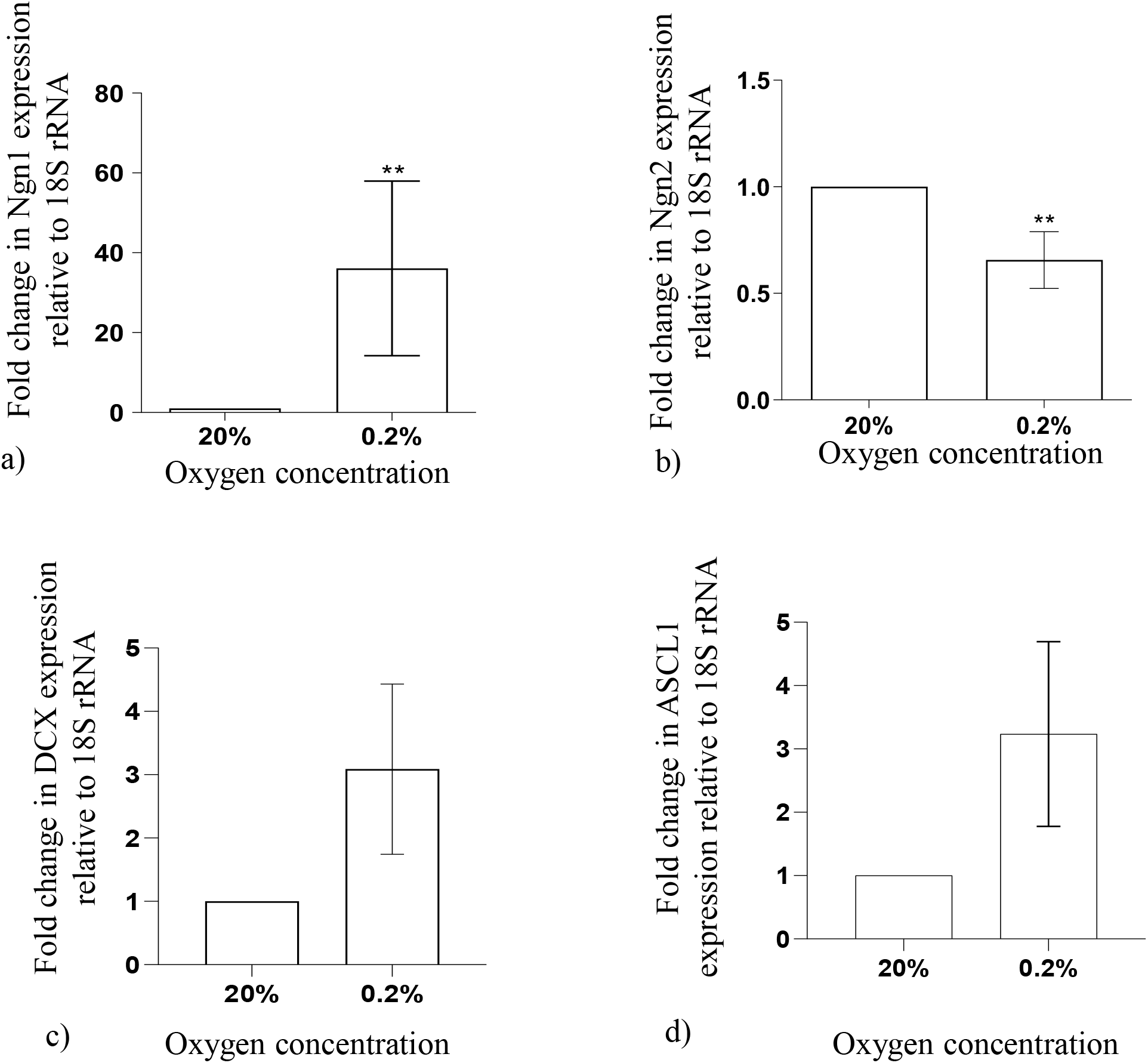
Expression of neuronal lineage markers, as analysed by qPCR (n=9), after exposure of human FNSCs to hypoxia (0.2% oxygen). a) Ngn1 b) Ngn2 c) DCX and d) ASCL1 gene expression. Data represented as Mean±SEM. **p-value <0.01.

**Figure 5.**
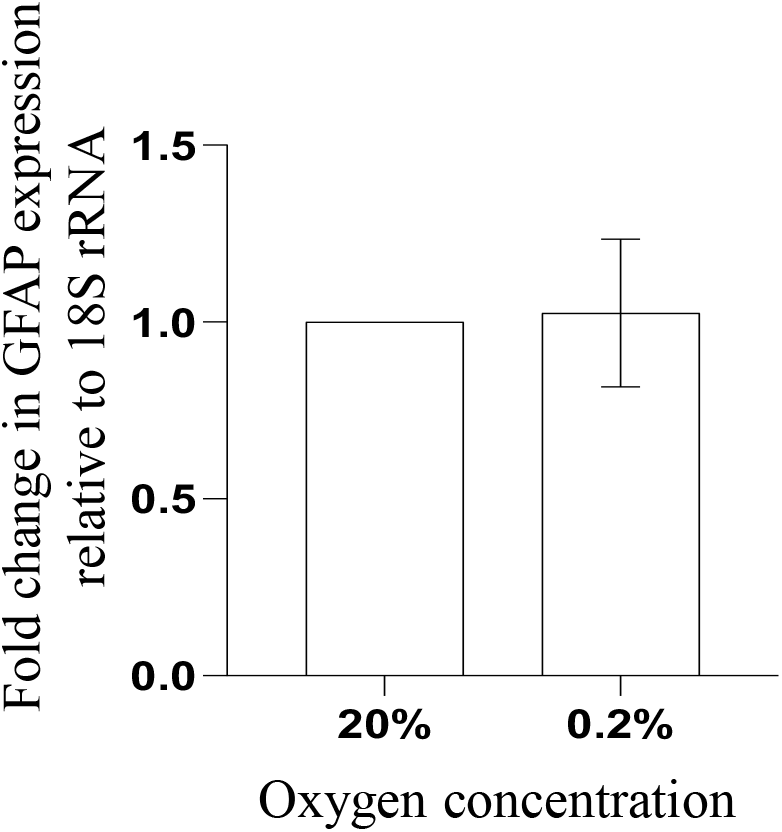
Expression of astrocyte lineage marker GFAP, as analysed by qPCR (n=9), after exposure of human FNSCs to hypoxia (0.2% oxygen). Data represented as Mean±SEM.

**Figure 6.**
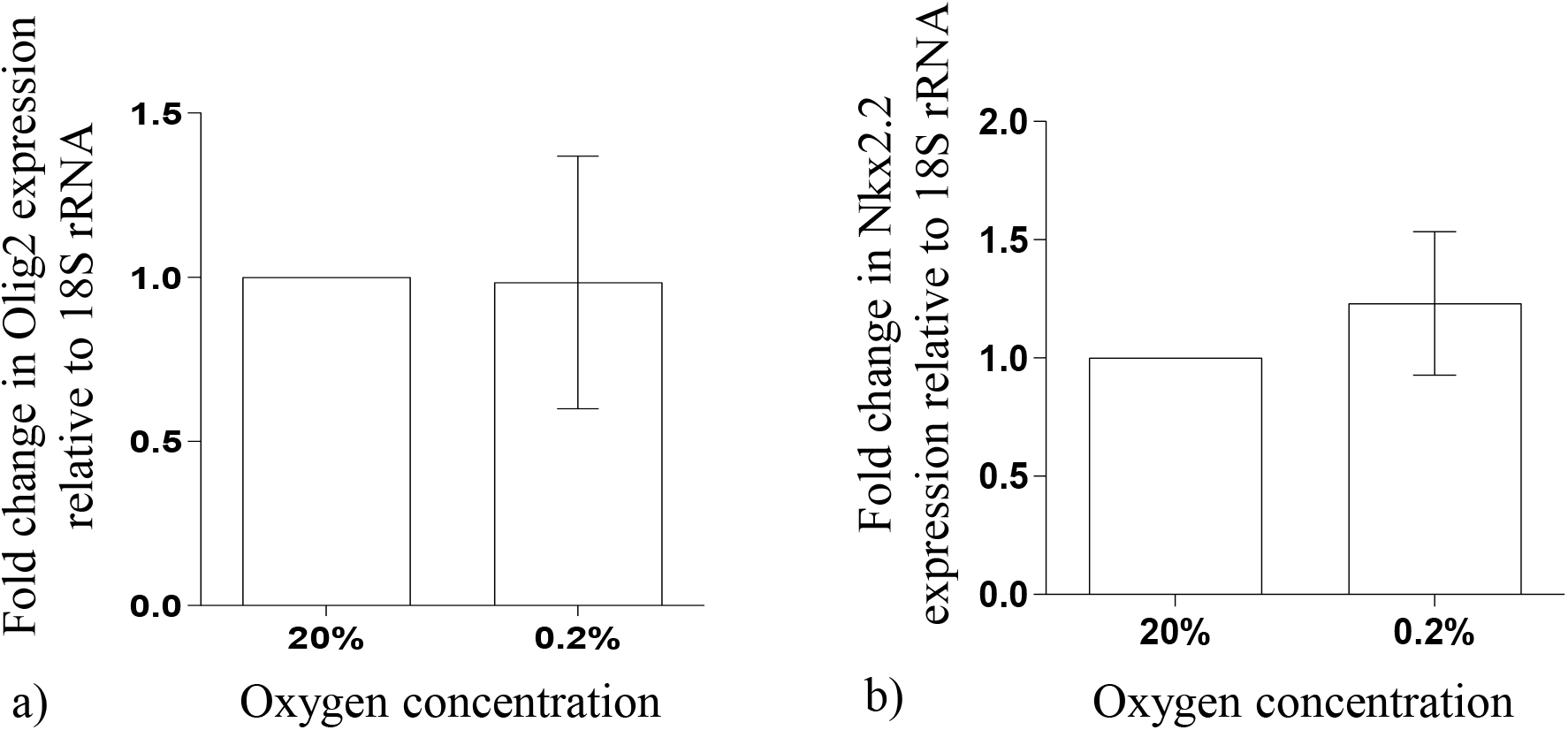
Expression of oligodendrocyte lineage markers, as analysed by qPCR (n=9), after exposure of human FNSCs to hypoxia (0.2% oxygen). a) Olig2 and b) Nkx2.2 gene expression. Data represented as Mean±SEM.

**Figure 7.**
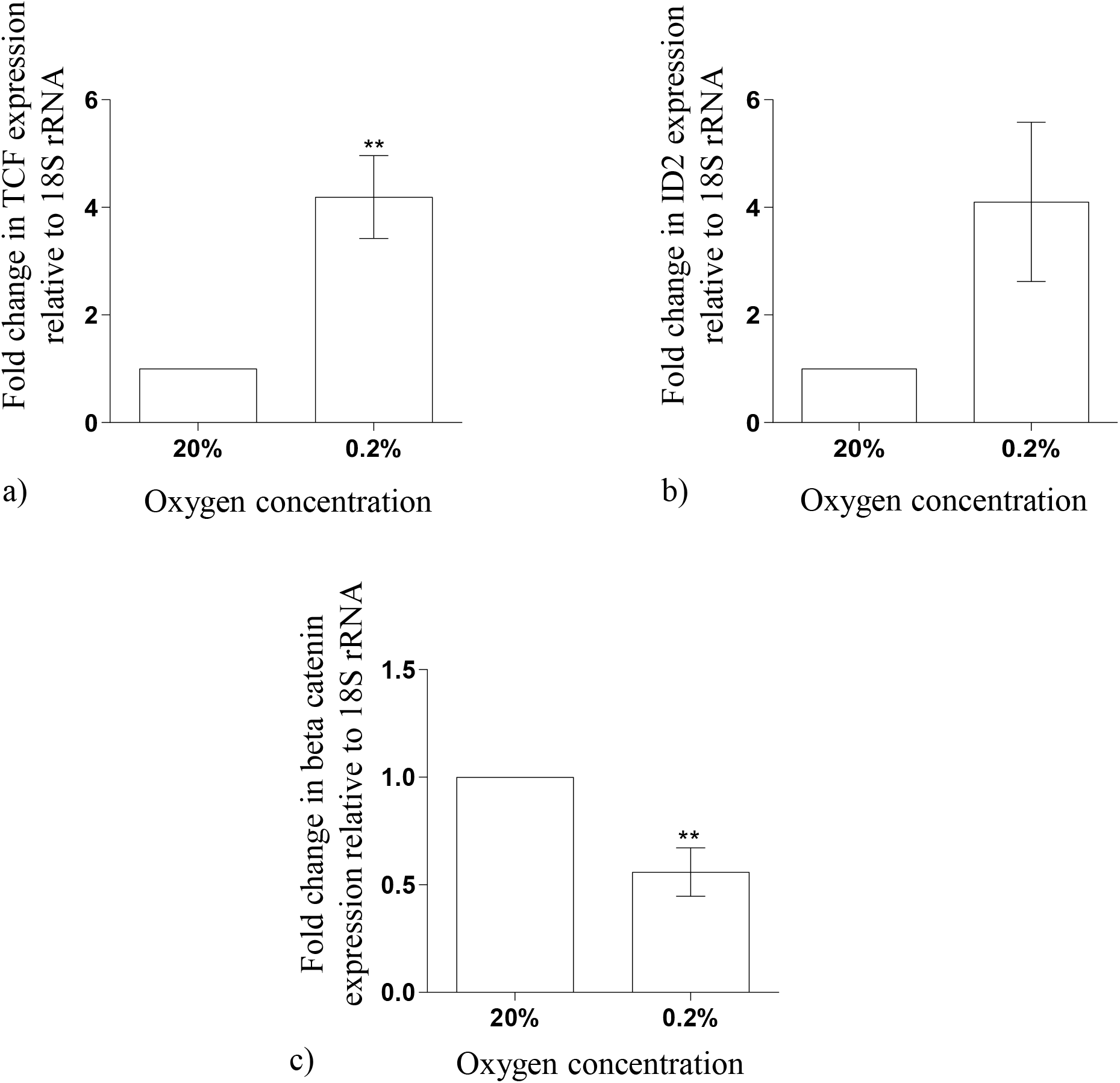
Expression of critical regulators of Wnt signaling pathway, as analysed by qPCR (n=9), after exposure of human FNSCs to hypoxia (0.2% oxygen). a) TCF b) ID2 and c) beta-catenin gene expression. Data represented as Mean±SEM. **p-value <0.01.

### Expression of genes involved in Wnt signaling pathway

Downstream targets of the Wnt signaling pathway beta catenin, TCF4 and ID2 were analyzed using qPCR. The mean fold change ± SEM for beta catenin, TCF4, and ID2 expression in the FNSCs exposed to hypoxia, were 0.55±0.11 (p=0.0426); 4.19± 0.77 (p=0.0352); and 4.10± 1.48 (p=0.2201) respectively.

## Discussion

In this study, human fetal neural stem cells (FNSCs) were isolated from subventricular zone of the aborted fetal brains. These multipotent stem cells, derived from neuroectoderm, have the potential to differentiate into neurons, astrocytes and oligodendrocytes. The isolated FNSCs displayed ability to form neurospheres, and expressed characteristic neural stem cell markers, Nestin and Sox2, as reported earlier (Bhagat *et al.*, 2018; Bernal and Arranz, 2018).

To mimic the physiological hypoxia *in utero*, human FNSCs were exposed to normoxia (20% oxygen) and hypoxia (0.2% oxygen), for 48 hours. Hypoxia mediates its action through HIF1α, and elevated levels of its downstream targets, carbonic anhydrase (CA9), phosphoglycerate kinase (PGK-1) and vascular endothelial growth factor (VEGF) indicate that the FNSCs were exposed to hypoxic environment. It also agrees with previous published reports related to downstream targets of HIF1α (Zhang, et al, 2018).

This study investigated the expression of early markers of neurogenesis like Ngn1, Ngn2, ASCL1 and DCX, in human FNSCs exposed to hypoxia. These are also considered to be the lineage commitment markers of neurogenesis. The expression of Ngn1, ASCL1 and DCX were found to be increased in human FNSCs exposed to hypoxia. However, the expression of Ngn2 was found to be slightly decreased. Interestingly, the increase in Ngn1 expression, was much more than the decrease in Ngn2 expression. This composite picture showing increased expression of Ngn1, ASCL1 and DCX signifies that hypoxia exposure in human FNSCs may be responsible for initiating neurogenesis, and thus promoting FNSCs to commit to neuronal lineage. Our findings are supported by reports indicating the essential role of Ngn1, ASCL1 and DCX in neurogenesis (Zhang et al, 2016; Pilz et al, 2018; Shahsavani et al, 2018). Interestingly, there are reports of different hypoxic conditions enhancing the expression of Ngn1 and DCX (Zhang et al, 2016; Jaworska et al, 2019; Haines et al, 2013). However, there are also a few studies reporting he downregulation of neurogenesis and ASCL1 by hypoxia, that are contrary to our findings (Vecera et al, 2020; Kasim et al, 2014). Again, studies also corroborate our findings by reporting that hypoxia or HIF-1α stimulates neurogenesis (Li et al, 2018; Varela-Nallar et al, 2014).

Our findings also indicate that the expression of GFAP, an astrocyte lineage commitment marker, and Olig2 and Nkx2.2, oligodendrocyte lineage commitment markers, were not influenced by exposing FNSCs to hypoxia. This indicates hypoxia not inciting FNSCs to commit to glial cell lineages. Our findings are corroborated by the fact that even though there are studies reporting reactive astrocytosis and gliosis in hypoxic injury, there are no reports that suggest promoting gliogenesis in neural stem cells (Zhang et al, 2017; Santilli et al, 2010).

This study used fetal neural stem cells as a model system to mimic fetal hypoxic conditions *in utero*. Analysis of whole genome transcriptomic changes and Gene ontology revealed that genes related to cell development and differentiation were differentially expressed in FNSCs exposed to hypoxia. On further analysis of these genes for biological processes, it was found that genes involved in neurogenesis were found to be up-regulated when FNSCs were exposed to hypoxia. Pathway analysis of our data indicated that the Wnt-beta-catenin signaling pathway may be implicated during this differentiation. This corroborates a previous report implicating this pathway in hypoxia mediated proliferation of neural stem cells (Qi et al, 2017).

Pathway analysis of microarray data displaying genes enriched in cell development and neurogenesis in our study, showed that canonical Wnt-beta catenin signaling pathway was involved in promoting commitment of FNSCs to neuronal lineage after exposing FNSCs to hypoxia. In this study, the expression of some critical regulators of Wnt-beta catenin signaling pathway, viz. beta catenin, TCF4 and ID2 were elucidated. Even though the expression of beta-catenin was slightly downregulated, the expression of its downstream effector targets, TCF4 and ID2, were increased in FNSCs exposed to hypoxia, indicating the involvement of the Wnt pathway in mediating the effects of hypoxia. As per previous reports, Wnt signaling is responsible for stem cell maintenance as well as differentiation, lineage commitment, axon guidance, and neurite outgrowth (Ille and Sommer, 2005). It has also been reported that Wnt signaling facilitates neurological recovery in experimental stroke, thereby establishing its role in neurogenesis (Song et al, 2019).

Beta catenin one of the regulators of the Wnt-signaling pathway, translocates to the nucleus and interacts with TCF to activate transcriptionally active complex. Interestingly, target genes of this complex are Ngn1 and Ngn2 (Kan et al., 2004). Ngn1/2 then binds to p300/CBP co-activator proteins to promote neuronal differentiation (Shin et al., 2016). The expression of TCF4 was found to be increased in this study, which might also explain the increase in the expression of Ngn1. This is probably the mechanism involved in influencing FNSCs to differentiate into neurons, after they are exposed to hypoxia. Recent studies have also confirmed the involvement of the Wnt-signaling pathway in mediating neurological recovery in stroke and epilepsy, lending credence to our findings in hypoxic conditions *in utero* (Sun et al., 2021; Song et al., 2019).

To the best of our knowledge, this is the first study to show that there was an increase in lineage commitment markers of neurogenesis viz., Ngn1, DCX and ASCL1, on exposing FNSCs to hypoxic conditions, while observing no change in the astrocytic and oligodendrocytic lineage markers. The mechanism attributed to the increased neurogenesis may be attributed to increase in downstream effectors of the Wnt-signaling pathway, viz. TCF4 and ID2 pointing to the involvement of the Wnt-signaling pathway in mediating this action. However, more functional experiments and analyses are needed to ascertain this.

## Acknowledgements

The authors are grateful to DBT, Govt. of India, for financial support – extramural research grant (BT/PR21413/MED/122/40/2016). The authors also wish to acknowledge the support of the facilities provided under the Biotechnology Information System Network (BTISNET) grant, DBT, Govt. of India, and Distributed Information Centre at NBRC, Manesar, India.

